# PIP3 depletion rescues myoblast fusion defects in human rhabdomyosarcoma cells

**DOI:** 10.1101/2020.03.06.980706

**Authors:** Yen-Ling Lian, Kuan-Wei Chen, Yu-Ting Chou, Ting-Ling Ke, Bi-Chang Chen, Yu-Chun Lin, Linyi Chen

## Abstract

Myoblast fusion is required for myotube formation during myogenesis, and defects in myoblast differentiation and fusion have been implicated in a number of diseases, including human rhabdomyosarcoma. While the transcriptional regulation of the myogenic program has been studied extensively, the mechanisms controlling myoblast fusion remain largely unknown. This study identified and characterized the dynamics of a distinct class of blebs, termed bubbling blebs, which are smaller than those that participate in migration. The formation of these bubbling blebs occurred during differentiation and decreased alongside a decline in phosphatidylinositol-(3,4,5)-trisphosphate (PIP3) at the plasma membrane before myoblast fusion. In a human rhabdomyosarcoma-derived (RD) cell line that exhibits strong blebbing dynamics and myoblast fusion defects, PIP3 was constitutively abundant on the membrane during myogenesis. Targeting phosphatase and tensin homolog (PTEN) to the plasma membrane reduced PIP3 levels, inhibited bubbling blebs, and rescued myoblast fusion defects in RD cells. These findings highlight the differential distribution and crucial role of PIP3 during myoblast fusion and reveal a novel mechanism underlying myogenesis defects in human rhabdomyosarcoma.

**Summary Statement:** This study reveals a novel mechanism underlying myogenesis defects in human rhabdomyosarcoma cells involving PIP3, whose depletion with PTEN rescues myoblast fusion defects.

## Introduction

Skeletal muscle is the most abundant tissue in vertebrates. Fusion of myoblasts is a decisive process for myotube formation. For muscular dystrophy/atrophy, recovery of muscle tissue highly depends on the differentiation and fusion of myoblasts to generate myofibers (Di Gioia et al., 2017). While the transcriptional mechanisms governing muscle development have been comprehensively studied in mammals (Moncaut et al., 2013), the cellular machinery underlying myoblast fusion remains unclear. Myoblast fusion involves several critical steps, including cadherin-mediated recognition, cytoskeleton-dependent adhesion, and membrane fusion (Beckett and Baylies, 2007; Kim et al., 2015). The muscle-specific membrane micropeptides, myomaker and myomixer (a.k.a. minion or myomerger), have been reported to regulate myoblast fusion (Bi et al., 2017; Millay et al., 2013; Zhang et al., 2017), with myomaker sufficient to induce fusion of non-muscle cells into myoblasts, which is enhanced by myomixer co-expression (Hernandez and Podbilewicz, 2017). Although numerous proteins and actin-myosin components have been implicated in myoblast recognition and fusion, the mechanisms driving morphogenesis of plasma membrane and lipid required for the fusion events remain largely unclear.

Previous studies investigating myoblast fusion in *Drosophila* have suggested that fusion competent cells generate podosome-like structures to “invade” founder cells before myofiber formation (Beckett and Baylies, 2007). Thus far, no fusion competent cells have been characterized in mammalian cells. Blebs are large membrane protrusions with roles in cell movement and invasion that can mark apoptotic cells (Charras, 2008; Sahai and Marshall, 2003). Plasma membrane protrusion is often caused by contraction of the actin cortex and the detachment of the membrane from the actin cytoskeleton (Tyson et al., 2014). Since phosphatidylinositol-4,5-bisphosphate (PIP2) controls the actin regulators and its depletion decreases membrane-cortex adhesion (Sheetz, 2001), PIP2 may be involved in bleb formation (Glaser et al., 1996; Hartwig et al., 1992).

In this study, we used time-lapse imaging to investigate a distinct class of small blebs, termed “bubbling blebs”, during differentiation of C2C12 and primary myoblast by comparing their dynamics, size, and motility to conventional blebs and determining their role in myoblast fusion alongside plasma membrane phospholipids. We also investigated whether manipulating phosphatidylinositol-3,4,5-trisphosphate (PIP3) levels in the plasma membrane could rescue differentiation defects associated with human rhabdomyosarcoma (RMS). Collectively, our findings reveal that bubbling blebs are involved in myoblast fusion and that PIP3 plays an important role in myogenesis and RMS.

## Results and Discussion

### Identification and characterization of bubbling blebs during myogenesis

Previous studies of amoebae, embryonic cells, and mammalian tumor cells have demonstrated that these cells can utilize blebs for migration as a common alternative to lamellipodia-driven migration (Friedl and Wolf, 2003). Skeletal muscle-derived C2C12 myoblasts and primary myoblasts were cultured in growth medium (GM), switched to low serum differentiation medium (DM) to induce myogenesis, and morphological change was monitored. Fusion events of differentiating myoblasts were systematically examined using time-lapse microscopy with 5-min intervals for 3 days. Differentiating myoblasts changed from a fibroblast-like morphology to a spindle-like morphology and their motility increased before fusion with other myoblasts or myotubes. Before fusion, 75% of the primary myoblasts displayed small membrane protrusions with an average diameter of 3.2 μm (Fig. 1A-1E), while C2C12 myoblasts displayed blebs with an average size of 4.3% of their maximum cell length, much smaller than previously described for satellite cells (7% cell length) (Otto et al., 2011). We termed these structures “bubbling blebs” and investigated their dynamics using lattice light sheet microscopy (Movie S1). Bubbling blebs were consistently distributed throughout the plasma membrane in differentiated, elongated, and mono-nucleated cells (Fig. 1B and Movie S2). Around 7-15% of myoblasts had a smooth cell surface with fewer, more polarized membrane protrusions (Fig. 1C-1E and Movie S2), thus were termed conventional blebbing cells (Fig. 1C). Primary myoblasts that displayed ambiguous bleb characteristics or no distinct features were categorized into “Unsure” (3%) or “No” (15%) categories, respectively (Fig. 1D), with quantitative analysis revealing that over 80% of myoblast fusion events displayed some forms of blebbing. These results suggest an important role of blebbing in myoblast fusion.

**Figure 1.**
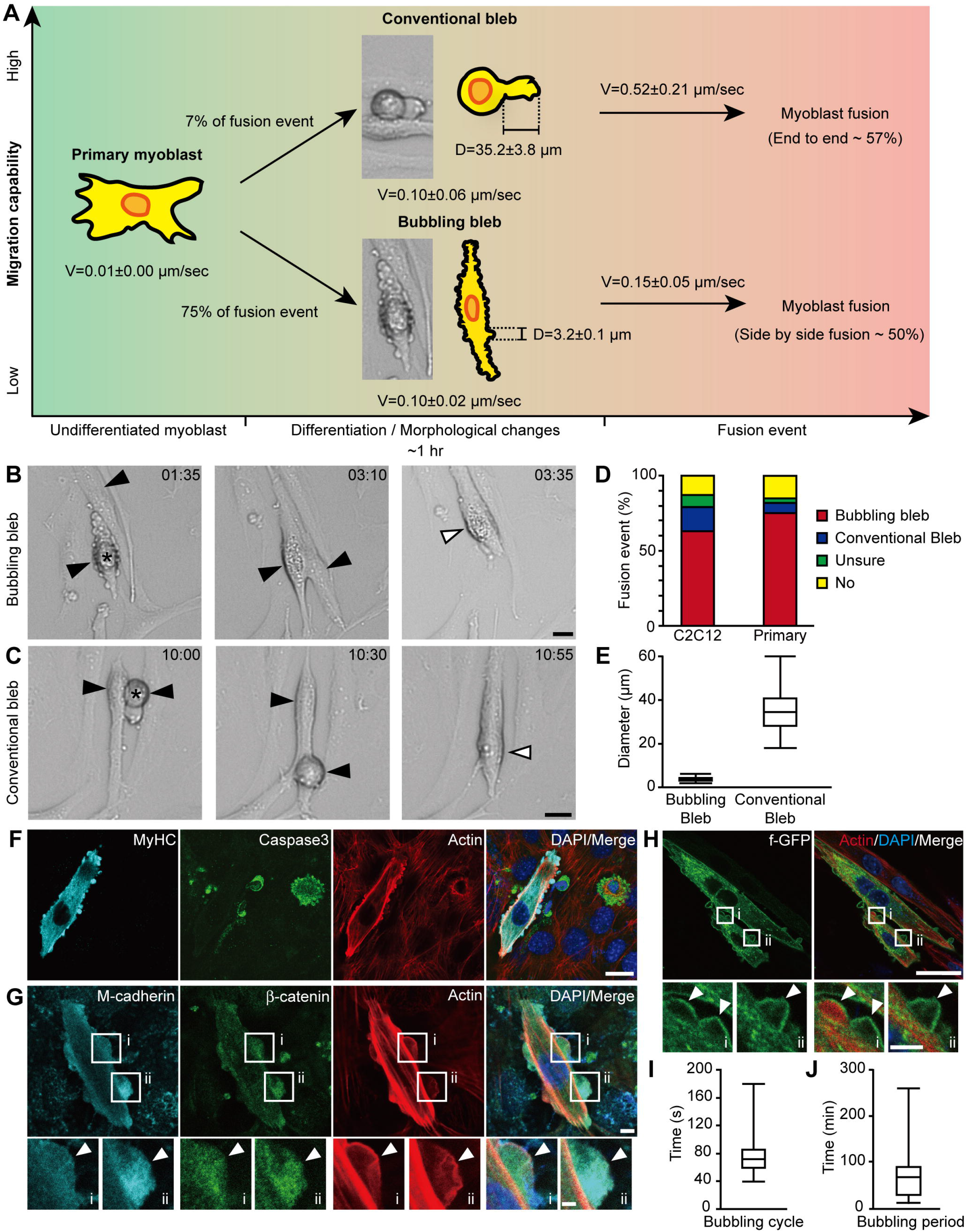
Identification and characterization of bubbling blebs during myogenesis. (A) Scheme of the distinct features of bubbling and conventional blebbing cells during myogenesis. (B-C) Differentiated primary myoblasts displaying bubbling or conventional bleb characteristics before fusion. Black arrowheads: fusion partners; white arrowheads: newly formed myotube (time, hr:min). (D) Relative fusion% for cells with bubbling, conventional blebs, cannot be determined (Unsure) or no (No) blebs before fusion. 200 and 60 fusion events were calculated for C2C12 cells and primary myoblasts, respectively. (E) Mean diameters of bubbling and conventional blebs are shown. (F-G) C2C12 cells were subjected to immunofluorescence staining at DM2d and followed by confocal microscopy. Arrowheads in (G) indicate the bubbling blebs. Anti-MyHC, anti-caspase 3 antibodies, rhodamine phalloidin and DAPI staining were shown in (F). Anti-M-cadherin, anti-β-catenin antibodies, rhodamine phalloidin and DAPI staining were shown in (G). (H) C2C12 cells were transfected with farnesylated GFP (f-GFP), differentiated and subjected to immunofluorescence staining with rhodamine phalloidin and DAPI. (I) Average cycle time for individual bubbling bleb for C2C12 cells is shown. (J) Average bubbling period for primary myoblasts during myogenesis. Box plots are from at least three independent experiments with more than 10 cells in (E) and 30 cells in (I-J). Scale bars are (B-C): 20 μm; (F, H): 10 μm and 2 μm for enlarged images in (H); (G): 5 μm and 2 μm for enlarged images.

To exclude the possibility that bubbling blebs were from apoptotic cells, differentiating C2C12 cells were subjected to immunofluorescence staining with myogenesis marker, MyHC, and active Caspase 3, a marker for apoptosis. C2C12 cells with bubbling blebs expressed MyHC but not active Caspase 3, indicating that these cells were not apoptotic (Fig. 1F). To determine whether bubbling blebs were involved in myoblast fusion, we subjected differentiated C2C12 cells to immunofluorescence staining with the cell-cell recognition markers M-cadherin and β-catenin (Charrasse et al., 2007). M-cadherin and β-catenin were enriched on bubbling blebs while actin lined their periphery (Fig. 1G), also visualized using farnesylated GFP (f-GFP; Fig. 1H). To characterize the dynamic behavior of the bubbling blebs, we calculated the duration of their cycle using time-lapse imaging. While C2C12 cells had an average formation cycle of around 70 sec (Fig. 1I), bubbling blebs were present on primary myoblasts for around 67 min (Fig. 1J), indicating continuous membrane remodeling before myoblast fusion.

Next, we categorized the orientation of cell-cell fusion as side-by-side, end-to-end, and other (Fig. 2A-2C). The probability of different fusion orientations was similar in C2C12 cells and primary myoblasts, with the probability of side-by-side fusion being over 50% (Fig. 2D) and more than 60% of primary myoblasts with bubbling blebs displaying fusion initiativity, defined as myoblasts actively fuse with their partners (Fig. 2E). Since the movement of blebbing cells in search for suitable fusion partners could reflect fusion efficiency, we analyzed their migration path and motility before, during, and after bubbling/conventional blebbing (Fig. 2F-2H). There was no significant difference in the motility or migration of the two cell types before or after bubbling and conventional blebbing; however, myoblasts were prone to lamellipodia-based migration prior and posterior to bubbling/blebbing (Fig. 2F-2H). During blebbing, cells with conventional blebbing moved longer distance compared to those with bubbling blebs (Fig. 2G). In terms of velocity, cells with conventional blebs move significantly faster than those with bubbling blebs after blebbing period (Fig. 2I). After blebbing, the cell surface became smooth and generally fused with the surrounding cells within 4 hr (Fig. 2J). To determine the correlation between bubbling or conventional blebbing and fusion, we measured the percentage of cells with these structures 4 hr before fusion, excluding blebbing cells undergoing cell division. The number of bubbling myoblasts decreased from 8.2 to 6.2%, whereas that of conventional blebbing cells increased from 7.6 to 9.9% before fusion (Fig. 2K). Taken together, these data suggest that bubbling blebs represent a distinct class of morphodynamic structures in C2C12 cells and primary myoblasts that precede myoblast fusion. Moreover, cells with bubbling blebs exhibit lower motility and strong initiativity for myoblast fusion, compared to those with conventional blebs. Since muscle tissue regeneration requires the recruitment and fusion of satellites cells to the site of lesion (Bentzinger et al., 2012), the identification of these bubbling blebs allows us to predict candidate myoblasts for fusion, which has not been possible until now.

**Figure 2.**
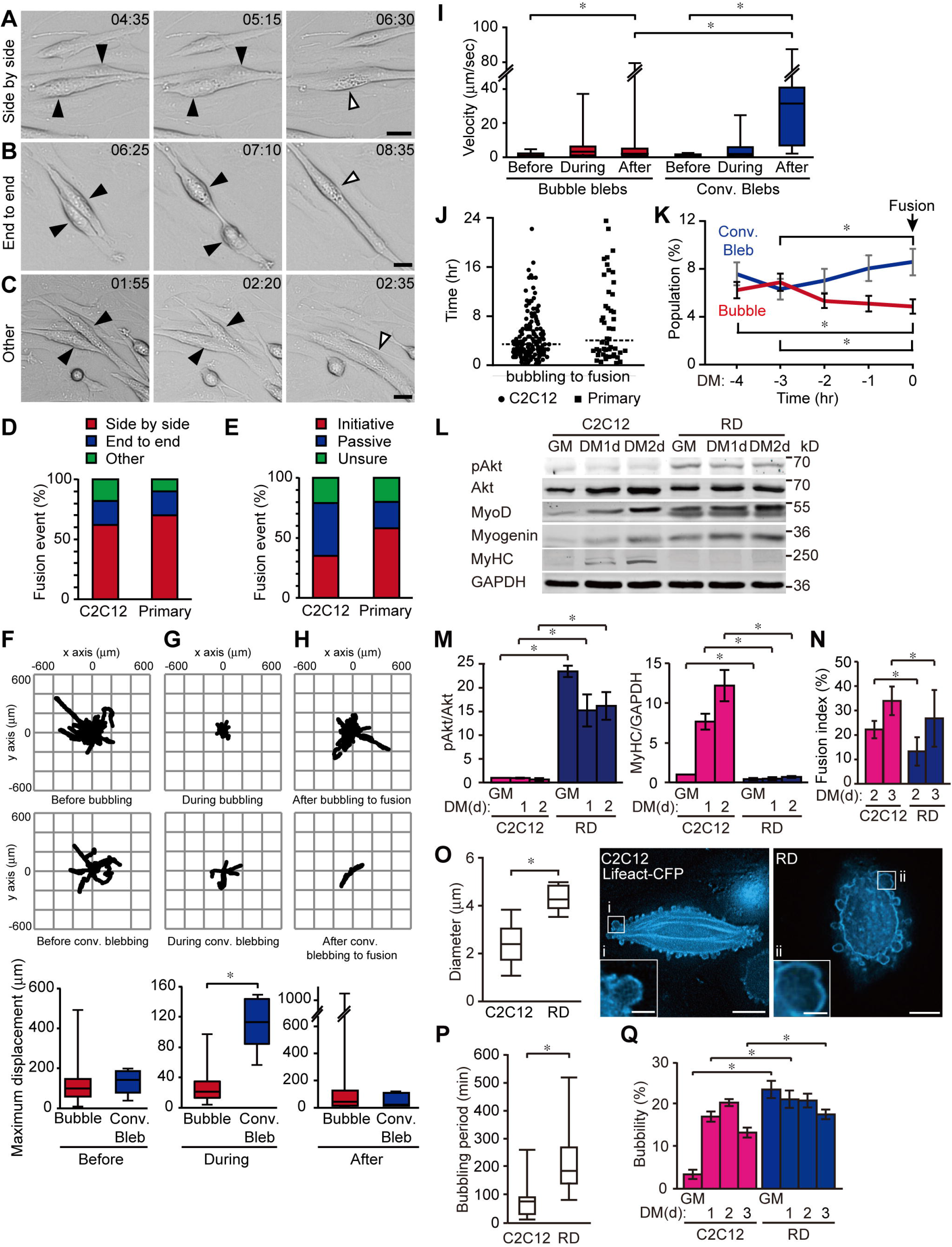
Correlation of bubbility and myoblast fusion for C2C12 and fusion-defective RD cells. (A-C) Primary myoblasts were differentiated and tracked individually by time-lapse microscopy with 5-min intervals for 3 days. Black and white arrowheads indicate the fusion partners and the newly formed myotube, respectively (time, hr:min). Categories of cell orientation before fusion: side-by-side, end-to-end or other. (D-E) Quantification of cell-cell fusion orientation and cell initiativity. Initiativity was defined as myoblasts actively fused to their partners. (F-H) Primary myoblast migration tracks before, during, and after bubbling or conventional bleb formation. Maximum displacement is the maximal distance that a cell moves. (I) Average velocity before, during, and after bubbling or conventional bleb formation as in (F-H). (J) Time period between cells displaying bubbling or conventional bleb characteristics and fusion. (K) Percentage of primary myoblasts displaying bubbling or conventional blebs 4 hr before fusion. 151 and 49 fusion events were calculated for C2C12 cells and primary myoblasts, respectively, in (D-K). (L) Indicated protein levels at GM and DM1d-2d in C2C12 and RD cells. (M) pAkt and MyHC levels were normalized to total Akt and GAPDH, respectively, then to C2C12 GM. (N) Fusion index in C2C12 and RD cells. (O) Left: mean diameter of bubbling blebs on C2C12 and RD cells. Right: cells transfected with Lifeact-CFP. (P) Bubbling period and (Q) bubbility (bubbling bleb formation) in C2C12 and RD cells. Values= mean ± s.e.m. and box plots are from at least three independent experiments with 10 cells calculated/experiment (O, P). At least three independent experiments were performed with more than 2200 cells/experiment in (N, Q) (\**P* < 0.05, one-way ANOVA for (F-H) and paired Student’s *t*-test for (I-Q)). Scale bars are (A-C): 20 μm; (O): 10 μm and 2 μm for enlarged images.

### Differentiation defects of RMS-derived (RD) cells

Human RMS, which is the most common soft tissue sarcoma in children under the age of ten, is presumed to originate from skeletal muscle due to its myogenic phenotype. There are two RMS subtypes, embryonal RMS (eRMS) and alveolar (aRMS). Histologically, eRMS resembles embryonic skeletal muscle, whereas aRMS is more aggressive and has a poorer prognosis. Although Pax3-Foxo1 or Pax7-Foxo1 translocation fusion is correlated with aRMS, the genetic profile of eRMS remains unclear (Kohsaka et al., 2014). In this study, we observed higher pAKT levels in eRMS-derived cells than in C2C12 cells (Fig. 2L-2M), potentially due to increased phosphoinositide-3-kinase (PI3K) activity or decreased phosphatase and tensin homolog (PTEN). In addition, RD cells expressed the myogenic markers MyoD and myogenin upon differentiation, like C2C12 cells; however, myosin heavy chain (MyHC) expression was inhibited in RD cells (Fig. 2L-2M) and the myoblast fusion index was significantly lower (Fig. 2N). Next, time-lapse imaging was employed to characterize the morphogenesis of RD cells during differentiation. Undifferentiated RD cells displayed similar bubbling blebs observed in differentiating C2C12 cells and primary myoblasts; however, bubbling blebs were significantly larger in RD cells, with an average size of 4.0 μm, compared to 2.5 μm in C2C12 cells (Fig. 2O). Furthermore, RD cells displayed a longer bubbling duration (Fig. 2P), with C2C12 cells exhibiting the highest bubbling activity at DM1d-2d during myogenesis but RD cells having a high bubbling bleb activity from GM until DM3d (Fig. 2Q). These results show a distinct morphodynamics during RD differentiation.

### PIP2, PIP3, and PS distribution in bubbling blebs

Since phospholipids and actin regulators are known to interact at the plasma membrane-cortex interface (Czech, 2000; Suetsugu et al., 2014), phospholipid distribution may affect morphogenesis and cell fusion during myogenesis. Phospholipid biosensors, pleckstrin homology (PH) domains of PLCδ (PHPLCδ), Akt (AktPH), and the C2 domain of Lactadherin (C2Lact), allow us to specifically detect PIP2, PIP3, and PS, respectively (Holz et al., 2000). To visualize the dynamics of these phospholipids within plasma membrane, C2C12 cells were transfected with lipid biosensors, differentiated, and followed by live-cell imaging (Movie S3). PIP2 signals were mainly found lining the protruded surface of the plasma membrane but not the base of the bubbling blebs (Fig. 3A, enlarged i panels). Conversely, PIP3 probe was distributed in the cytosol during differentiation and the apical surface of bubbling blebs, while PS signals were found on the protruded surface and basal side of the bubbling blebs (Fig. 3A, ii-iii-iv). Comparing the levels of PIP2, PIP3, and PS at GM and DM2d stages revealed that plasma membrane PIP3 levels declined during myogenesis (Fig. S1A-S1B).

**Figure 3.**
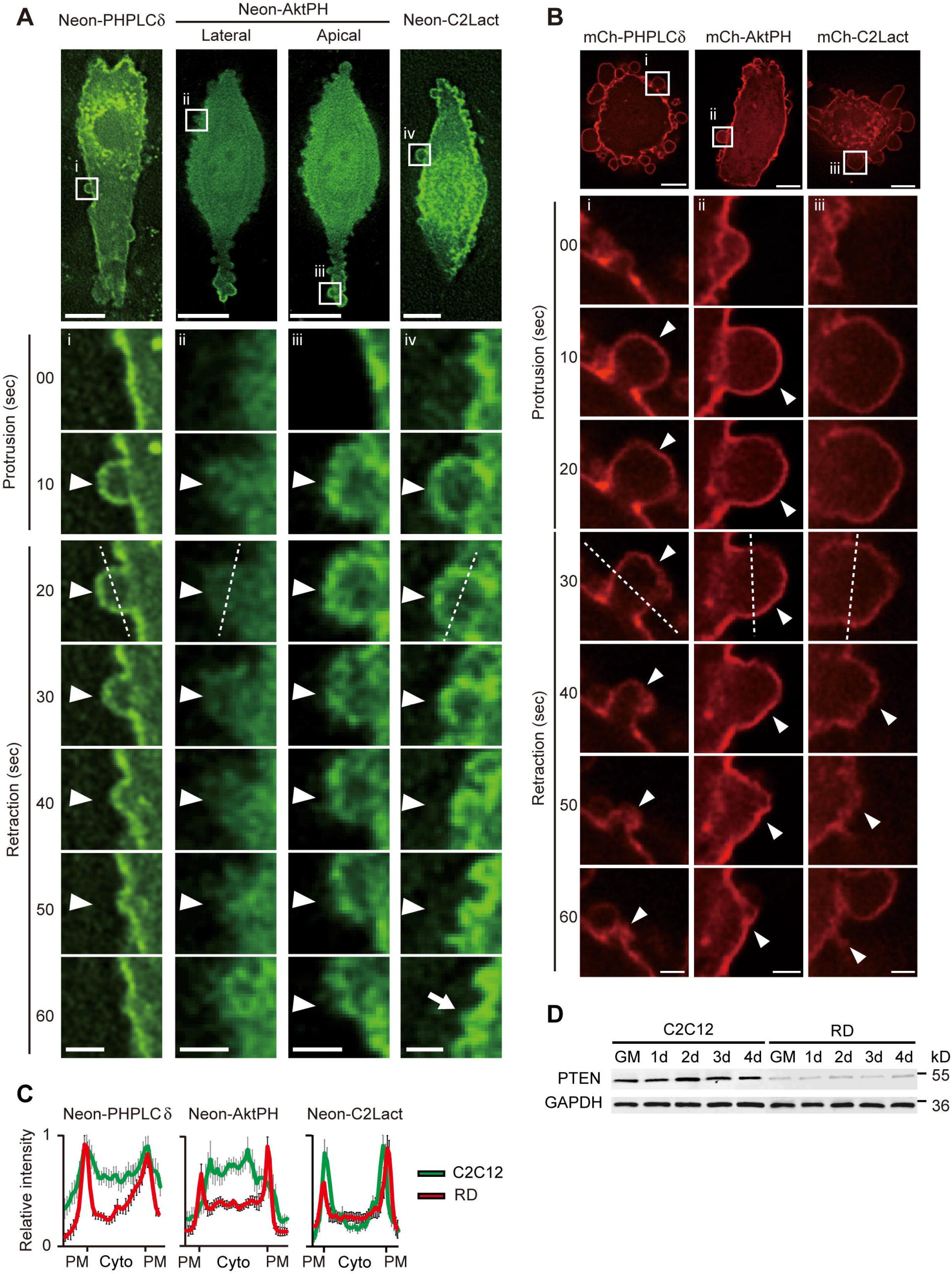
Role of PIP2, PIP3 and PS in bubbling bleb formation. Time-lapse imaging of differentiated C2C12 (A) and RD (B) cells transfected with fluorescently tagged (Neon or mCherry) phospholipid biosensors (-PHPLCδ: PIP2; -AktPH: PIP3; -C2Lact: PS) showing lateral (A-ii) and apical (A-iii) side close-ups and individually tracked bubbles (arrowhead). (C) Intensity histograms of PIP2, PIP3, and PS distribution on the plasma membrane (PM) and cytosol (Cyto) at 20 sec in C2C12 cells (green) and 30 sec in RD cells (red). Values= mean ± s.e.m. of 10-25 bubbling blebs. (D) PTEN protein levels in undifferentiated (GM) C2C12 and RD cells and at differentiation days 1-4 (1d-4d). Scale bars: 10 and 2 μm for enlarged images.

Next, we examined whether defect in PIP3 redistribution may contribute toward the pathogenesis of human RMS. For RD cells, the distribution of PIP2 and PS within the bubbles (Fig. 3B, i, iii) was similar to those in C2C12 cells (Fig. 3A, i, iv), whereas PIP3 probe was detected throughout the cell membrane in RD cells during differentiation (Fig. 3B, ii, and Movie S4), but mostly in the cytosol of differentiated C2C12 cells (Fig. 3A, ii). The relative fluorescence across the white dash lines in Fig. 3A-3B indicated the expression patterns of these PIs in C2C12 and RD cells (Fig. 3C). This finding reveals a differential distribution of PIP3 upon differentiation of C2C12 and RD cells, which may underlie the fusion defect of RD cells. One mechanism to remove PIP3 from cell surface is through de-phosphorylation of PIP3 to PIP2 by PTEN. Interestingly, PTEN levels were dramatically lower in RD cells than in C2C12 cells (Fig. 3D), consistent with previous findings (Zhu et al., 2016). Thus, these results suggest that PIP3 is redistributed during bubbling bleb formation and before fusion (Fig. S1), with its failure to deplete from the plasma membrane partly contributing to fusion defects in RD cells.

### Reducing PIP3 by targeting PTEN to the cell surface improves myoblast fusion

PTEN dephosphorylates PIP3 to PIP2 and thus antagonizes the action of PI3K to inhibit the AKT signaling pathway. To assess the role of PIP3 in bubbling bleb dynamics and myoblast fusion, chemically induced dimerization (CID) systems were applied, in which rapamycin induced the dimerization of FK506 binding protein (FKBP) and FKBP–rapamycin-binding domain (FRB; Fig. 4A) (Putyrski and Schultz, 2012). Membrane-targeting domain of LYN kinase was fused to CFP-FRB fusion protein, allowing Lyn-CFP-FRB anchoring to the plasma membrane (Inoue et al., 2005). Live-cell imaging of RD cells transfected with Lyn-CFP-FRB, YFP-FKBP, and mCherry-AktPH revealed that rapamycin-induced YFP-FKBP translocation had no significant effect on bubbling blebs in RD cells (Fig. 4B and Movie S5). However, rapamycin induced the translocation of YFP-FKBP-PTEN from the cytoplasm to the plasma membrane, reducing PIP3 signals at the plasma membrane (Fig. 4C and Movie S6) as shown by line-intensity histograms analysis (Fig. 4D). The velocity map generated using ADAPT revealed that RD bubbling bleb microdomains underwent continuous membrane expansion (green) and retraction (red) (Fig. 4E, top panel) which stopped with 54 min of PIP3 depletion as indicated as yellow regions (Fig. 4E, bottom panel). To examine whether PIP3 depletion on the plasma membrane would rescue myogenesis defect of RD cells, RD cells overexpressed Lyn-CFP-PTEN was differentiated for 1 or 2 days, fixed and performed immunofluorescence staining with anti-MyHC antibody and DAPI. PTEN overexpression on the plasma membrane had no significant effect on bubbility and fusion index, compared to control groups (Lyn-CFP), for C2C12 cells; however, bubbility was markedly reduced 36% (Fig. 4F) and fusion index was increased by 58% (Fig. 4G) for RD cells. These results are consistent with our finding that bubbling blebs preceded fusion and that their activity decreased immediately prior to fusion.

**Figure 4.**
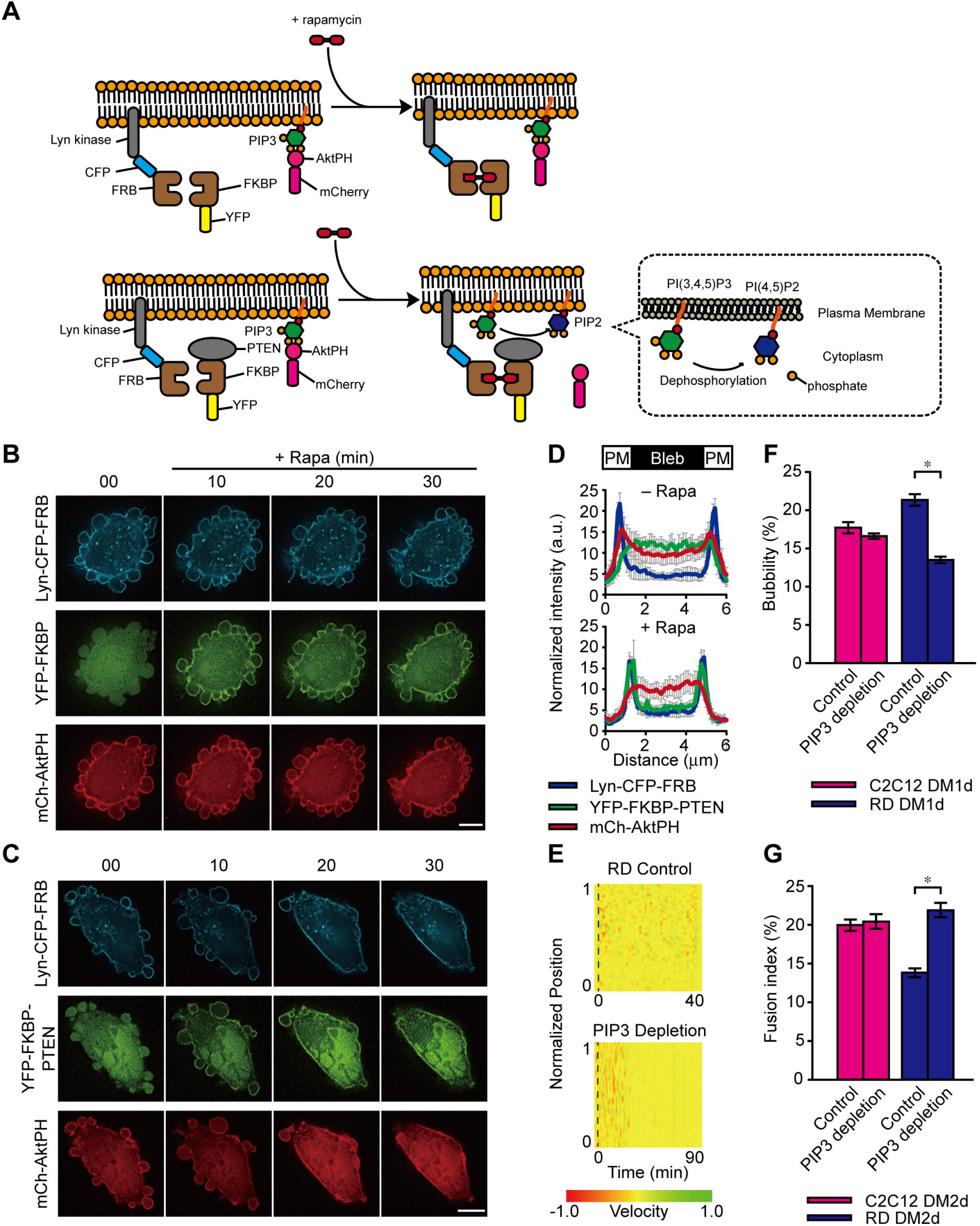
Targeting PTEN to the cell surface reduces PIP3 and improves myoblast fusion in RD cells. (A) Chemically induced dimerization (CID) system. Cells co-transfected with Lyn-CFP-FRB and mCh-AktPH, along with YFP-FKBP (top panel) or YFP-FKBP-PTEN (bottom panel), were treated with rapamycin to induce YFP-FKBP or YFP-FKBP-PTEN translocation to the Lyn-CFP-FRB-labeled plasma membrane. PTEN translocation induced PIP3 dephosphorylation (bottom panel). Time-lapse imaging at the indicated time points after rapamycin (100 nM) addition. (D) Intensity histogram of (C) demonstrating membrane anchor (blue), PTEN (green), and PIP3 (red) distribution on the plasma membrane and within bubbling blebs. (E) Velocity analysis of cells in (B-C) using ADAPT. Representative velocity map showing velocity changes in membrane protrusion (green) and retraction (red) over time in RD (top panel) and PIP3-depleted RD (bottom panel) cells. Black dash lines indicate rapamycin treatment time. (F-G) C2C12 and RD cells were transfected with Lyn-CFP-PTEN, differentiated and subjected to immunofluorescence staining with anti-MyHC antibody and DAPI. Quantification of bubbility at DM1d (F) and fusion index at DM2d (G). Values= mean ± s.e.m. of three independent experiments with 23-28 bubbling blebs in RD cells (D), and more than 2200 cells/experiment in (F, G) (\**P* < 0.05, paired Student’s *t*-test). Scale bars: 10 μm.

In conclusion, these findings suggest that bubbling bleb formation primes myoblast fusion, possibly via plasma membrane remodeling and PIP3 redistribution, and reveal a possible mechanism underlying myogenesis defects in human RMS. Moreover, reduced PTEN levels in RD cells stabilize PIP3 residence at the cell surface and constitutively activate AKT signaling which hampers myoblast fusion, while temporal PIP3 depletion at the plasma membrane and reduced bubbling bleb formation are required to facilitate RD cell fusion.

## Materials and Methods

### Reagents

Dulbecco’s Modified Eagle Medium (DMEM), fetal bovine serum (FBS), horse serum (HS), L-glutamine (L-Gln), antibiotic-antimycotic (AA), penicillin-streptomycin (P/S), Lipofectamine 2000, 6-diamidino-2-phenylindole (DAPI), ProLong Gold, rhodamine phalloidin, Hoechst-33342, Alexa Fluor 700 goat anti-mouse, 647 goat anti-mouse and 488 donkey anti-rabbit IgG secondary antibody were purchased from Invitrogen (Carlsbad, CA). IRDye800CW-labeled anti-rabbit secondary antibody was purchased from LI-COR Bioscience (Lincoln, NE). F-10 medium and pronase were purchased from Sigma-Aldrich (St. Louis, MO). Fibroblast growth factor 2 (FGF2) was purchased from ProSpec (Rehovot, Israel). Bovine serum albumin (BSA) and bicinchoninic acid (BCA) was purchased from Santa Cruz Biotechnology (Santa Cruz, CA), and TransIT-LT1 was purchased from Mirus (Madison, WI). The following antibodies were used: anti-MyHC and anti-Myogenin (Developmental Studies Hybridoma Bank, #Cat MF20, 1:100 for IF and 1:1000 for WB; #Cat F5D, 1:1000 for WB, respectively); anti-M-cadherin (BD Transduction Laboratories™, #Cat 611100, 1:100 for IF); anti-β-catenin (Millipore, #Cat 06-734, 1:100 for IF); anti-Caspase 3, anti-Akt and anti-phospho-Akt (Ser473) and anti-non-phosphorylated PTEN (Cell Signaling Technology, #Cat 9661, 1:100 for IF; #Cat 9272, 1:1000 for WB; #Cat 4051, 1:2000 for WB; #Cat 7960, 1:1000 for WB, respectively); anti-MyoD (C-20) (Santa Cruz Biotechnology, #Cat SC-304, 1:1000 for WB); GAPDH (GeneTex, #Cat GTX627408, 1:5000 for WB). (IF: Immunofluorescence staining; WB: Western Blotting)

### Cell culture, animal handling and ethics statement

C2C12 cells (60083) and RD cells (60113) were purchased from Bioresource Collection and Research Center of Food Industry Research and Development Institute, Taiwan, and maintained at sub-confluent densities in growth medium (DMEM supplemented with 10% FBS, 1% AA, and 1% L-Gln) at 37°C and 5% CO_2_. Upon reaching confluency, the cells were switched to differentiation medium (DMEM supplemented with 2% HS, 1% AA, and 1% L-Gln) to induce myoblast fusion.

Primary myoblasts were isolated as described previously (Bondesen et al., 2004). All animal experiments were under conduct in according to the guidelines of the Laboratory Animal Center of National Tsing Hua University (NTHU), Taiwan. Animal usage protocols were reviewed and approved by the NTHU Institutional Animal Care and Use Committee (Approval number 10607). Briefly, 12- to 14- week-old Sprague Dawley female adult rats were sacrificed and their tibialis anterior muscle was dissociated and washed with PBS followed by DMEM with 1% P/S and 1% L-Gln. The muscles were minced into small pieces in DMEM containing 1% P/S and 1% L-Gln, digested with 0.1% pronase for 1 h at 37°C, and centrifuged at 2000 rpm for 3 min. The mixture was re-suspended in DMEM containing 10% FBS, 1% P/S, and 1% L-Gln, and then filtered using a 70 μm nylon mesh. Single isolated primary myoblasts were collected, re-suspended in F-10 medium with 5 ng/mL of FGF2 and 0.5 μg/mL of heparin, and then cultured on collagen-coated dishes at 37°C and 5% CO_2_. Primary myoblasts were switched to the same differentiation medium to induce fusion.

### Plasmids, transfection and viral infection

Farnesylated GFP (f-GFP) was a gift from Professor Chuan-Chin Chiao at the National Tsing Hua University. Lifeact-CFP, mCherry-PHPLCδ, mCherry-AktPH, mCherry-C2Lact, Neon-AktPH, Lyn-CFP, Lyn-CFP-FRB and YFP-FKBP-PTEN were constructed in previous study (Fan et al., 2017). mCherry-PHPLCδ was subjected to PCR to provide cleavage sites at 5’ and 3’ ends of PHPLCδ domain (forward: 5’- CCGCTCGAGCTATGGACTCGGGCCGGGAC-3’ and reverse 5’-CCGGAATTCTCTAGACTGGATGTTGAGCTCCTTCAG-3’). The PCR product was inserted into Neon.c1 plasmid to generate Neon-PHPLCδ, while mCherry-C2Lact was sub-cloned into Neon.c1 to generate Neon-C2Lact. FKBP-YFP-PTEN was subjected to PCR with cleavage sites at the 5’ and 3’ ends of full-length PTEN (forward: 5’-CATGGACGAGCTGTACAA-3’ and reverse: 5’-AAGGAGCTCCGGTGGATCCTCAGACTTT-3’). The PCR product was inserted into the Lyn-CFP-FRB.c1 plasmid to generate Lyn-CFP-PTEN. C2C12 cells were transiently transfected with the plasmids of interest using Lipofectamine 2000 or TransIT-LT1 according to the manufacturer’s instructions. After 24 to 48 hr, the cells were differentiated and processed by immunofluorescence staining or time-lapse imaging.

### Fluorescence microscopy, time-lapse imaging, and image data analysis

Cells were fixed in 4% paraformaldehyde, permeabilized by 0.1% Triton X-100, and subjected to immunofluorescence staining with the indicated antibodies and dyes. Immunofluorescence images were detected by Carl Zeiss Observer Z1 microscope and Carl Zeiss LSM 780 confocal microscope. Time-lapse microscopy was performed using a Carl Zeiss Observer Z1 and a Nikon Eclipse Ti inverted microscope, equipped with a cell incubator maintaining 5% CO_2_ at 37°C. Images were analyzed by AxioVision Rel. 4.8 and NIS-Elements Advanced Research. For some images, deconvolution software was applied to remove out-of-focus information using Huygens Essential. Images and composite figures were prepared using Adobe Photoshop and Illustrator. Relative fusion event was calculated based on whether differentiating myoblasts with bubbling or conventional blebs showed initiativity, defined as actively moving toward their stationary fusion partners. Passive fusion events were defined as bubbling or conventional blebs did not initiate the fusion events. Cells displaying ambiguous fusion initiativity or passive fusion characteristics were placed in the “unsure” group. Movement of individual migrating cells was analyzed by manual tracking in AxioVision software. The center of a cell was identified and selected in each frame. The relative x-y positions were then created as tracks in the Microsoft Excel. Cell movement was segmented based on three different time periods: before, during and after bubbling/conventional blebs. Maximum displacement is the maximal distance of a cell moves from the zero point. The average velocity of the cells was calculated by dividing the maximum displacement by the corresponding time period.

Cells containing at least three nuclei were considered as myotubes. The fusion index (%) was calculated as the number of nuclei (≥3) in myotubes divided by total number of nuclei in MyHC-positive cells (indicating differentiating cells). Among all the MyHC-positive cells in each frame, bubbility was calculated as the percentage of cells with bubbling or conventional blebs. ADAPT (v1.185) software was used as an online plug-in for ImageJ according to the developer’s instructions (Barry et al., 2015). ADAPT segmented each individual frame of the cytosolic channel using a region-growing algorithm, with the seed point for the frames determined automatically. The cell boundary contour was taken as the set of foreground points bordering the background in each segmented frame, resulting in a time-varying contour. By adjusting the threshold algorithms available in ImageJ, individual bubbling blebs in the time-lapse movies were tracked automatically, exported as velocity map and the mean bubbling velocity was analyzed.

### Western blotting

Cells were harvested by radioimmunoprecipitation assay (RIPA) buffer containing 1 mM phenylmethylsulfonyl fluoride (PMSF), 1 mM Na_3_VO_4_, 10 ng/ml aprotinin, and 10 ng/ml leupeptin (A+L). Protein samples were determined by BCA assays and an equal amount of proteins from each sample was separated by sodium dodecyl sulfate-polyacrylamide gel electrophoresis (SDS-PAGE), followed by incubation with the indicated primary antibodies and IRDye-conjugated secondary antibodies. Signal was detected using an Odyssey infrared imaging system (LICOR Biosciences).

### Statistical analysis

Histological analyses and parametric data were presented as the mean ± s.e.m. Differences between groups were evaluated using paired Student’s *t*-tests, whereas multiple comparisons were analyzed by one-way ANOVA with repeated measurement. *P* values of < 0.05 were considered statistically significant.

## Competing interests

The authors declare no competing or financial interests.

## Author contributions

Conceptualization: LC, YCL, YLL, KWC; Methodology: YLL, KWC, BCC, YCL; Validation: YLL, YTC, TLK; Formal analysis: YLL, YTC, TLK; Investigation: YLL, LC, YCL; Resources: LC, YCL; Data curation: YLL, KWC, BCC, YTC, TLK; Writing - original draft: LC, YCL, YLL; Writing - review & editing: LC, YLL, YCL, TLK, KWC; Visualization: YLL, YCL, YTC, TLK; Supervision: LC, YCL; Project administration: LC and YCL; Funding acquisition: LC, YCL.

## Funding

This work was supported by the Ministry of Science and Technology, Taiwan (Grant # MOST 108-2320-B-007-005-MY3 to LC; 107-2628-B-007-001, 108-2636-B-007-003, 109-2636-B-007-003, and 108-2638-B-010-001-MY2 to YCL) and by the National Health Research Institutes, Taiwan (Grant# NHRI-EX109-10813NI to LC).

